# Warm temperature and mild water stress cooperatively promote root elongation

**DOI:** 10.1101/2023.11.30.569400

**Authors:** Scott Hayes, Cheuk Ka Leong, Wenyan Zhang, Marthe Lamain, Jasper Lamers, Thijs de Zeeuw, Francel Verstappen, Andreas Hiltbrunner, Christa Testerink

**Affiliations:** Laboratory of Plant Physiology, Wageningen University, Wageningen 6700 AA, The Netherlands; Institute of Biology II, Faculty of Biology, University of Freiburg, Freiburg, Germany; Signalling Research Centres BIOSS and CIBSS, University of Freiburg, Germany

**Author notes:** Email address for correspondence: Scott Hayes and Christa Testerink.

## Abstract

Warm temperatures have a dramatic effect on plant development. In shoots, stems elongate, and leaves are raised in a developmental programme called thermomorphogenesis. This results in enhanced leaf cooling capacity^1^. Thermomorphogenesis is tightly intertwined with light signalling pathways. The level of integration is so high that it has been proposed that shoot temperature sensing may have evolved from the co-option of an existing light signalling pathway during the colonisation of land by plants^2^. Roots also undergo thermomorphogenesis, but the mechanism by which this occurs is less well understood. Main root elongation is enhanced at warm temperatures, and this response is independent of many of the light and temperature signalling components of the shoot^3^. Roots develop in darkness and so it is a reasonable assumption that root temperature signalling is not through modulation of light signalling. It was recently speculated that due to the close correlation between warm temperature and soil moisture content, root temperature signalling could feasibly be related to water availability signals^2^. In this study we tested the interaction between temperature and water availability signalling in plant roots. We found that these environmental factors co-operatively enhance main root elongation. This interaction effect was dependent on SUCROSE NON-FERMENTING RELATED KINASE 2.2 (SnRK2.2) and SnRK2.3 and the E3 ubiquitin ligase CONSTITUTIVELY PHOTOMORPHOGENIC 1 (COP1). We found that SnRK2.2 / 2.3 and COP1 have opposite effects on the stability of the transcription factor ELONGATED HYPOCOTYL 5 (HY5) in elongation zone hair cells. The stability of HY5 in these cell types generally corresponded to the degree of root elongation seen in each mutant background. Our study reveals several molecular components of root thermomorphogenesis and highlights the importance of an integrative approach to plant environmental signalling. Our results may have direct implications for agricultural land management, especially as global climates become more unpredictable.

## Results and Discussion

To investigate the interaction between temperature and water availability on root growth, we grew Arabidopsis seedlings on soil with a range of water contents, at either 20°C or 28°C. Water content was re-established daily through watering the soil surface. We found when the soil water content was high (23% to 19% water by weight-approx. 107% to 89% of field capacity), warm temperature did not induce root elongation (Figure 1A). Only at soil water contents of 17% (approx. 79% field capacity) or lower did we observe root thermomorphogenesis (Figure 1A). We noticed that these plants had a slight hypocotyl phenotype. In well-watered soil, warm temperature strongly promoted hypocotyl elongation, as described previously^4^. This effect tended to decrease somewhat in drier soils albeit not to a statistically significant degree (Figure 1B). This effect is reminiscent of the recently reported inhibition of shade-induced hypocotyl elongation under drought^5^. We compared the root and hypocotyl lengths of each plant in our assay and found a weak negative correlation at 28°C, but not at 20°C (Figure S1A-B). This correlation did however appear to be mostly a product of drought treatment groups, rather than any within treatment effect. SnRK2 kinases are well-established regulators of water stress signalling in plants^6^ and so we tested whether these genes are required for the interaction between warm temperature and water stress on Arabidopsis seedling architecture. We repeated our experiment at 21% and 17% soil water contents in the wild type and *snrk2*.*2 / snrk2*.*3* mutant. We found that wild type plants showed mild temperature-induced root elongation at 21% soil water content, but that the effect of temperature was greatly enhanced at 17% soil water content (Figure 1C-D). The roots of a *snrk2*.*2 / snrk2*.*3* mutant however behaved similarly to the wild type at 21% soil water content and did not show a synergistic effect of warm temperature and water availability (Figure 1C-D). Warm temperature-induced hypocotyl elongation was slightly reduced by mild water stress in wild type plants, but this was less pronounced in mutants lacking SnRK2.2 and SnRK2.3 (Figure 1E).

**Figure 1.**
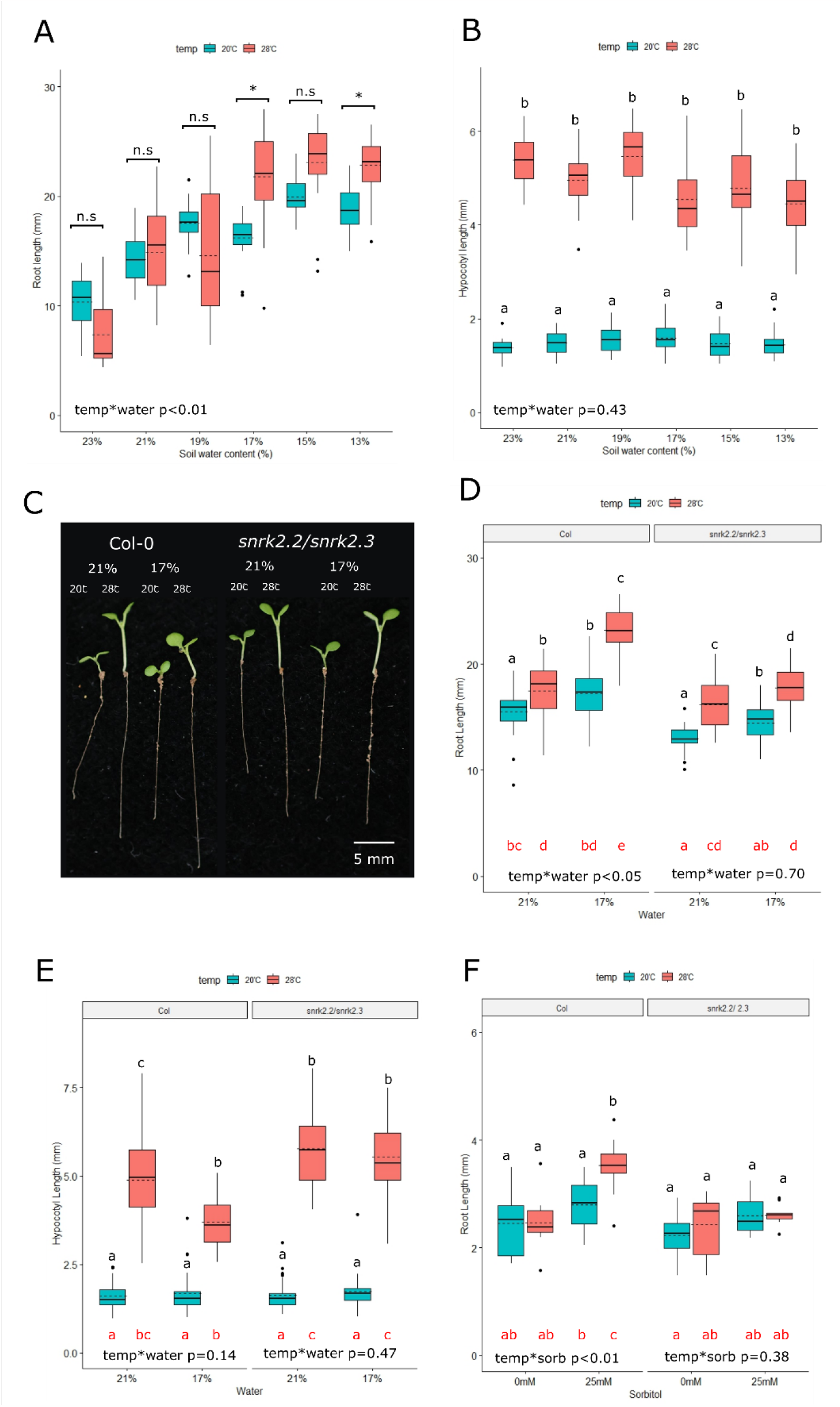
Warm temperature and mild water stress co-operatively promote root elongation. **A)** Root length of wild type (Col) arabidopsis plants grown on soil at different temperatures and water availabilities. Plants were sown directly onto soil of the indicated water content and germinated for 3 days under a propagator lid (150μE white light, 16h photoperiod, 20°C). Pots were then moved to 20°C or 28°C with the same lighting conditions for a further 4 days. Soil water content was re-established daily (n ≥ 19). **B)** Hypocotyl lengths of the plants grown in A (n ≥ 19). **C)** Representative images of Col and *snrk2*.*2* / *snrk2*.*3* plants grown at 21% and 17% water as in A. **D)** Root lengths plants grown as in C (n=30). **E)** Hypocotyl lengths of the plants grown in D (n=30). **F)** Root lengths of Col and *snrk2*.*2* / *snrk2*.*3* seedlings that were germinated for 3 days in white light (150μE, 16h photoperiod, 20°C) on 1/10^th^ MS media, before roots were detached and placed on 1/10^th^ MS media supplemented with 5 mM sucrose and 0 mM or 25 mM sorbitol (n=14). Roots were grown in darkness for another 4 days at either 20°C or 28°C before measurement. In each panel, *within genotype* interaction terms between environmental conditions are shown. In panels A, B, D and E, a Scheirer Ray Hare test and Dunn’s post hoc test with a Benjamini-Hochberg correction were used. In panels F and G, a 2-way ANOVA with Tukey’s HSD test was used. Black letters or asterisks indicate statistically significant means (p< 0.05) within each genotype. Red letters indicate statistically significant means (p< 0.05) across the whole experiment.

Roots can autonomously respond to warm temperature treatments^3^, but it has also been shown that there is a genetic linkage between shoot and root thermomorphogenesis^7^. To investigate whether our root phenotype was root autonomous, we adopted a detached root assay^3^. We first grew detached roots on plates in the dark with a range of supplementary sucrose concentrations (Figure S1C). In the absence of sucrose, we did not observe any root growth. At 5mM sucrose, we observed reasonable root elongation, but not thermomorphogenesis. As sucrose concentrations increased however, roots gained the capacity to respond to temperature. Sucrose acts as both an osmolyte and energy source. To investigate whether the increase in osmotic stress was driving thermomorphogenesis, we grew detached roots on 5mM sucrose, supplemented with a range of sorbitol concentrations (Figure S1D). We found that sorbitol also promoted root thermomorphogenesis, and that this was dependent on SnRK2.2 and/ or SnRK2.3 (Figure 1F). We were also able to induce the capacity for thermomorphogenesis in Arabidopsis by supplementing roots with mannitol or sodium chloride at equimolar concentrations (Figure S1E). We observed a similar pattern of responses in lettuce, leek, and radish roots, albeit with a weaker interaction term. This raises the possibility that water availability and warm temperature cooperatively induce root elongation in a variety of angiosperms (Figure S1F-H).

SnRK2s are critical positive regulators of abscisic acid (ABA) signalling^8^. ABA is involved in a large number of stress responses. We therefore tested whether ABA could also confer roots the capacity to elongate at warm temperature. We found that low concentrations of ABA promoted root thermomorphogenesis, and that this was also dependent on SnRK2 kinases (Figure 2A). Severe osmotic stress is known to promote the accumulation of ABA^8^ and so we next tested whether ABA signalling was induced by mild osmotic stress and warm temperature. To this end we utilised the *6xABRE-A:erGFP* reporter line, that contains six copies of the abscisic acid response element from *ABA INSENSITIVE 1 (ABI1)*, driving the expression of an endoplasmic reticulum localised GFP^9^. Surprisingly, we could not detect any effect of sorbitol on GFP fluorescence in the root tip, instead we saw an increase in GFP signal at warm temperatures, particularly in the QC and columella stem cells (Figure 2B, S2A-B). We were however unable to detect any changes in ABA levels (in whole roots) across our experimental conditions (Figure 2C). We also observed an interaction between osmotic stress and warm temperatures on root elongation in several mutants that are severely deficient in ABA signalling (Figure 2D-E). We therefore propose that although ABA can mimic the effect of osmotic stress on warm temperature-induced root elongation (Figure 1F and Figure 2A), it is not activation of the canonical ABA signalling pathway *per se* that drives the combined response.

**Figure 2.**
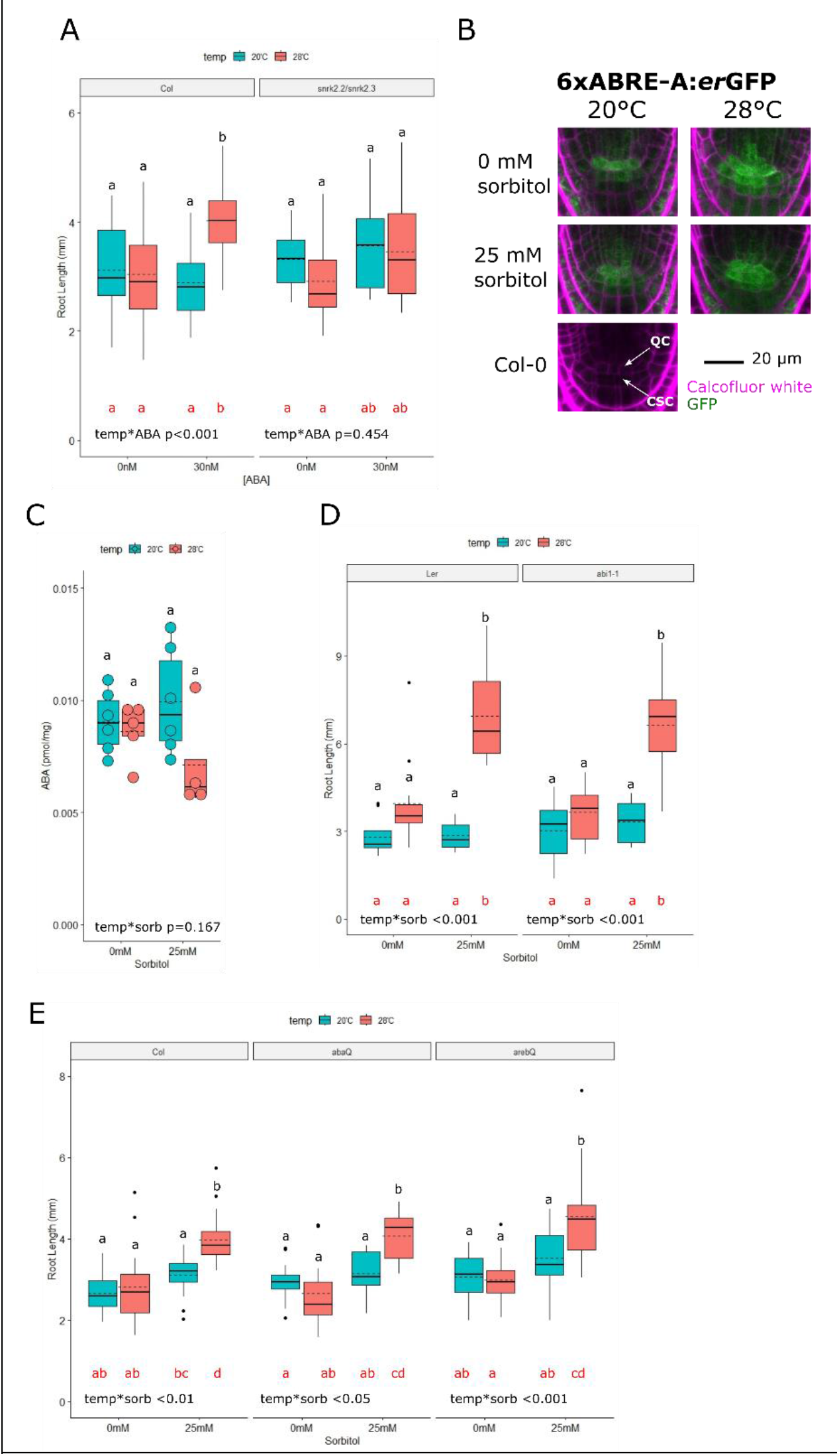
Warm temperature promotes ABA signalling in the root tip. **A)** Root lengths of Col and *snrk2*.*2* / *snrk2*.*3* seedlings that were germinated for 3 days in white light (150μE, 16h photoperiod, 20°C) on 1/10^th^ MS media, before roots were detached and placed on 1/10^th^ MS media supplemented with 5 mM sucrose and 0 nM or 30 nM ABA. Roots were grown in darkness for another 4 days at either 20°C or 28°C before measurement (n=16). **B)** Representative sum-stack images of the root tips of a 6xABRE-A:*er*GFP reporter line and Col wild type control. Plants were germinated for 3 days in white light (150μE, 16h photoperiod, 20°C) on 1/10^th^ MS media, before roots were detached and placed on 1/10^th^ MS media supplemented with 5 mM sucrose and 0 mM or 25 mM sorbitol. Roots were grown in darkness for another 4 days at either 20°C or 28°C, before fixation. Quiescent centre (QC) and Columella stem cells (CSC) are indicated. **C)** ABA levels in the roots of wild type Col plants grown as in B (n ≥ 4). **D)** Root lengths of wild type Ler and abi1-1 plants grown as in B (n=12). **E)** Root lengths of wild type Col, abaQ (*pyr1-1/pyl1-1/pyl2-1/pyl4-1*) and arebQ (*areb1/ areb2/ abf3/ abf1-1*) mutants grown as in B (n=16). In each panel, *within genotype* interaction terms between environmental conditions are shown. A 2-way ANOVA with Tukey’s HSD test was performed. Black letters indicate statistically significant means (p< 0.05) within each genotype. Red letters indicate statistically significant means (p< 0.05) across the whole experiment.

In an effort to establish a molecular pathway for the control of mild osmotic stress and warm temperature-induced root elongation, we tested mutants of genes that have established roles in shoot temperature signalling. The detached roots of plants deficient in *PIF1, 3, 4, 5* and *7* (*pifq/pif7-1*) behaved very similarly to the wild type (Figure 3SA), confirming early reports that these genes are not required for local root responses to warm temperature^10,11^. We also found that the roots of mutants lacking phyA and phyB were very similar to wild type roots (Figure S3B). *COP1* is essential for increased hypocotyl elongation at warm temperature^12^. We tested whether *COP1* plays a similar role in promoting root elongation. Surprisingly, we found that the *cop1-4* mutant has very high root elongation at warm temperatures, both in the presence and absence of osmotic stress (Figure 3A). The *cop1-6* mutant showed a similar trend, to a lower extent (consistent with the weak phenotype of this mutant in the dark^13^). Light represses COP1 activity^14^ and so we tested the effect of light on our assay. Indeed, we found that light conferred roots with the ability to respond to warm temperature, even in the absence of sorbitol (Figure 3B). We observed no additional effect of light in *cop1-4* mutant (Figure 3SB). These results reveal COP1 to be a repressor of warm temperature / water stress signalling in roots. Most studies into warm temperature-induced root elongation are performed in the light^3,7,10,15,16^. This result could therefore explain why multiple groups have reported warm temperature-induced elongation in the absence of water stress, and why warm temperature-enhanced root elongation is reduced or lost in short photoperiod conditions^15,16^.

**Figure 3.**
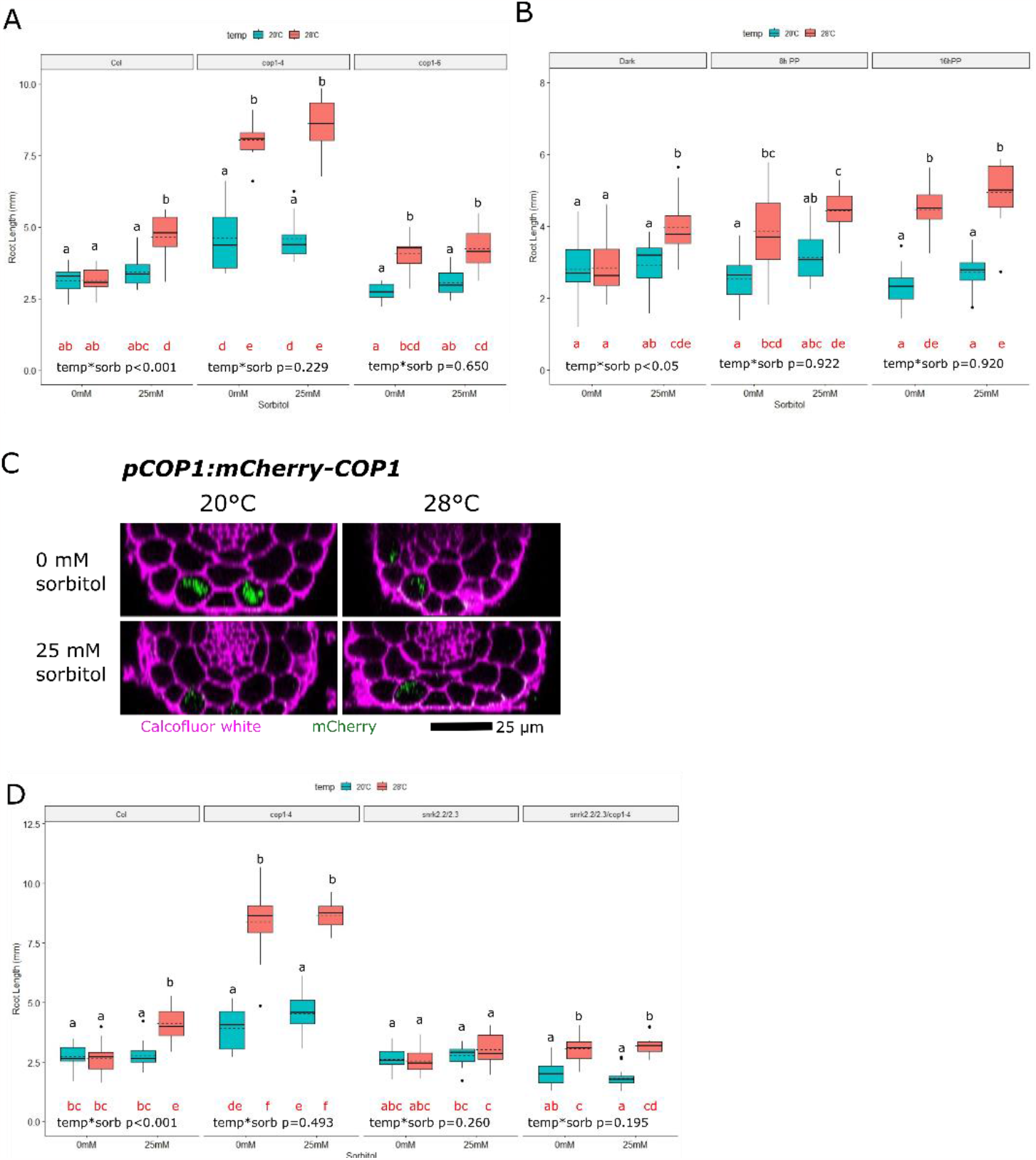
COP1 supresses warm temperature signalling in the absence of osmotic stress. Wild type Col, *cop1-4* and *cop1-6* mutant plants were germinated for 3 days in white light (150μE, 16h photoperiod, 20°C) on 1/10th MS media, before roots were detached and placed on 1/10th MS media supplemented with 5 mM sucrose and 0 mM or 25 mM sorbitol. Roots were grown in darkness for another 4 days at either 20°C or 28°C before being measured (n ≥ 8). **B)** Wild type Col plants were grown as in A but grown for the final 4 days in either darkness, 8 hours photoperiod (8h PP) or 16h PP white light (150μE) (n ≥ 13). **C)** Representative images of *pCOP1:mCherry-COP1* reporter lines grown as in A. Orthogonal sections were generated from the elongation zone. **D)** Root lengths of wild type Col, *cop1-4, snrk2*.*2/snrk2*.*3* and *cop1-4 / snrk2*.*2/ snrk2*.*3* lines grown as in A (n=16). In each panel, *within genotype* interaction terms between environmental conditions are shown. In panels A and D, a 2-way ANOVA with Tukey’s HSD test was performed. In panel B a Scheirer Ray Hare test and Dunn’s post hoc test with a Benjamini-Hochberg correction was used. Black letters indicate statistically significant means (p< 0.05) within each genotype. Red letters indicate statistically significant means (p< 0.05) across the whole experiment.

We next questioned whether warm temperature or osmotic stress influenced COP1 abundance in the root. To this end, we produced a line expressing *mCherry-COP1* under its native promotor (*pCOP1:mCherry-COP1*). We were able to detect mCherry fluorescence in only a few specific cell types, namely the epidermal hair cells of the elongation zone and the endodermis of the differentiation zone and mature root (Figure S3C). When we grew these plants in our experimental conditions, we found that the mCherry signal in epidermal hair cells was highest at 20°C in the absence of sorbitol. Warm temperature and the presence of sorbitol both reduced the mCherry signal in this cell type (Figure 3C, S3D-E). This suggests that COP1 abundance is reduced at either 28°C or in the presence of mild osmotic stress. Our initial hypothesis was that SnRK2s could contribute to root elongation in our experimental conditions through the suppression of COP1. However, we did not see a reduction in mCherry-COP1 abundance upon the application of ABA (Figure S3D-E). In fact, we saw that the application of ABA increased the abundance of mCherry-COP1 at 28°C. We also crossed our reporter into the *snrk2*.*2 / snrk2*.*3* background to see how this would affect mCherry-COP1 abundance. We found that in the *snrk* mutant background, nuclear mCherry-COP1 was barely detectable, lending support for the argument that the SnRKs are required to maintain COP1 abundance (Figure S3D-E). We found this result puzzling, as mutants with low COP1 abundance showed enhanced warm temperature-induced root elongation even in the absence of sorbitol (Figure 2A). We therefore tested the genetic interaction between COP1 and the SnRKs by generating *cop1-4/snrk2*.*2/2*.*3* triple mutants. As previously, the *cop1-4* mutant elongated in response to temperature with and without mild osmotic stress, and the *snkr2*.*2 / snrk2*.*3* mutant had lower root elongation than the wild type (Figure 3D). In the triple mutant however, we observed a similar pattern of root elongation as in the *cop1-4* mutant, but with a greatly reduced amplitude (Figure 3D). This suggests that enhanced root elongation in the absence of COP1 requires the presence of SnRKs. It also shows that warm temperature can promote root elongation in the absence of SnRK2.2 / SnRK2.3 and COP1.

To gain more insight into the regulatory network controlling root elongation in mild osmotic stress and warm temperature, we focused on identifying factors that could act downstream of COP1 and the SnRKs. HY5 was recently identified as a key regulator of plant root thermomorphogenesis^15^. COP1 is known to regulate HY5 abundance in response to warm temperature in the shoot^12^ and so we tested whether this regulatory module plays a role in the root. Indeed, we found that the exaggerated root elongation of the *cop1-4* mutant was highly dependent on the presence of *HY5* (Figure 4A). HY5 has previously been shown to be stabilised by warm temperature in the root ^15^. We therefore tested how HY5 stability is regulated in combined warm temperature and mild osmotic stress conditions, using a line expressing GFP-HY5 under a constitutive promoter^17^. Because we observed COP1 in only a subset of epidermal cells in the elongation zone (Figure 3C), we first investigated whether HY5 stability was differentially regulated in hair and non-hair cells. We found that in non-hair cells (where mCherry-COP1 was not detected) warm temperature promoted the stability of HY5 in both the presence and absence of sorbitol (Figure S4A-B). In hair cells by contrast, we found that warm temperature reduces HY5 stability in absence of sorbitol but promotes HY5 stability in the presence of sorbitol (Figure S4A, Figure 4B-C). We crossed the HY5 reporter line into the *cop1-4* background. We found that in the absence of COP1, hair cell localised HY5 is stabilised by warm temperature in both be presence and absence of sorbitol (Figure 4B-C). This pattern is similar to that of root elongation in the *cop1-4* mutant (Figure 4A), suggesting that the misregulation of HY5 may be responsible for enhanced root elongation in this mutant. We also crossed our GFP-HY5 reporter into the *snrk2*.*2 / snrk2*.*3* background. We found that the stability of HY5 in hair cells was dramatically reduced in the absence of *SnRK2s* (Figure 4B-C, Figure S4A). Interestingly though, despite its low abundance, the pattern of HY5 regulation was comparable to the original reporter line. This suggests that SnRK2s are important for maintaining HY5 abundance, but that warm temperature and mild osmotic stress are still able to regulate HY5 abundance in the absence of SnRK2.2 and SnRK2.3.

**Figure 4:**
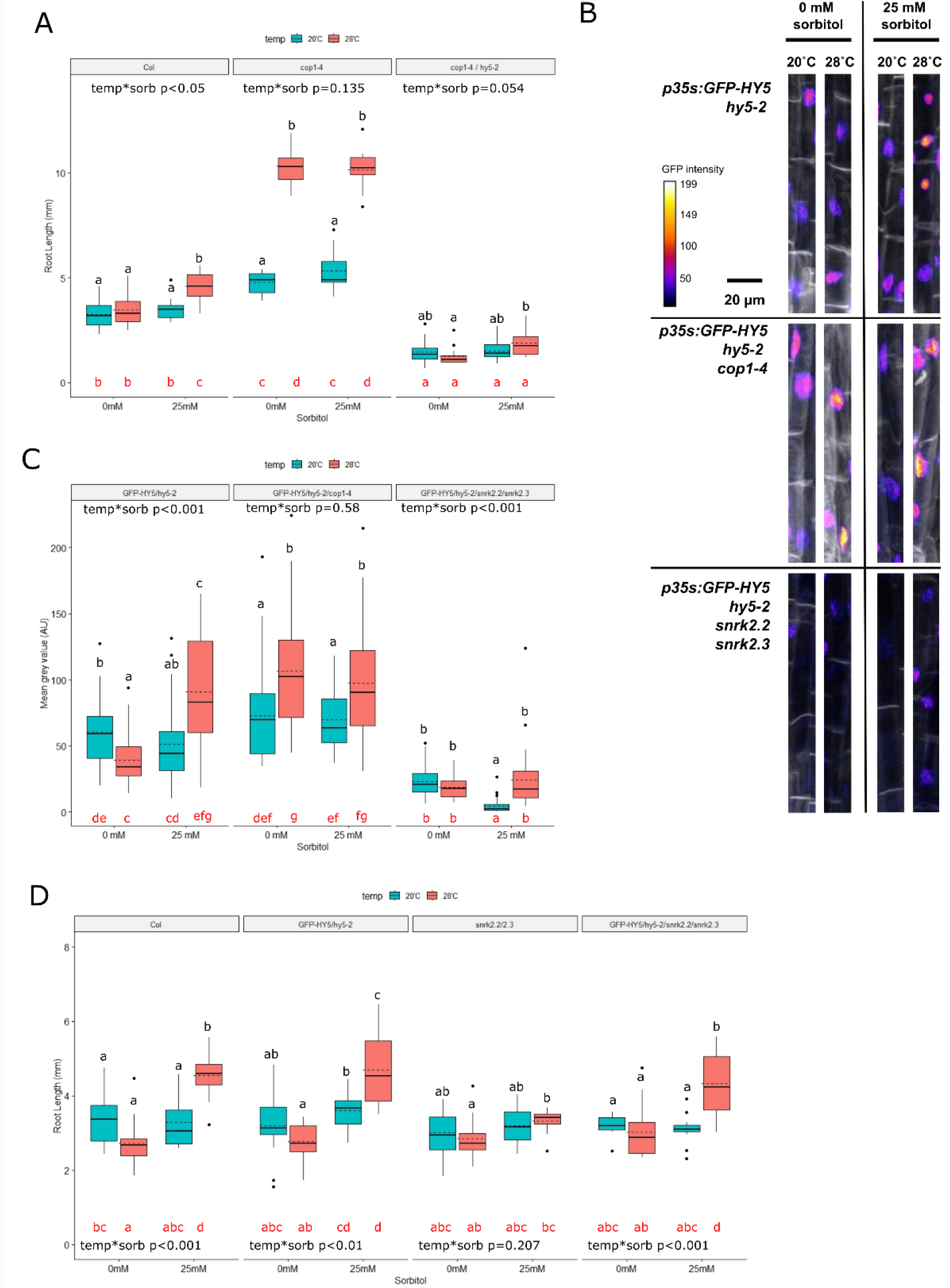
Warm temperature and mild water stress promote the accumulation of HY5. **A)** Root lengths of Col, *cop1* and *cop1* / *hy5-2* mutants grown for 3 days in white light (150μE, 16h photoperiod, 20°C) on 1/10^th^ MS media, before roots were detached and placed on 1/10^th^ MS media supplemented with 5 mM sucrose and 0 mM or 25 mM sorbitol. Roots were grown in darkness for another 4 days at either 20°C or 28°C before measurement (n=14). **B)** Sum slice image of the elongation zone hair cells of a *p35S:GFP-HY5* reporter in the *hy5, hy5*/*cop1* and *hy5* / *snrk2*.*2* / *snrk2*.*3* backgrounds. Seedlings were grown as in A before fixation. Calcofluor white stained cell walls are shown in grey. **C)** Mean grey value of hair cell nuclei from the images generated in B. Five nuclei were quantified from each root (n ≥ 6 roots). **D)** Root lengths of Col, *p35S:GFP-HY5 / hy5-2, snrk2*.*2* / *snrk2*.*3* and *p35S:GFP-HY5 / hy5-2* / *snrk2*.*2* / *snrk2*.*3* mutants grown as in A (n=14) In each panel, *within genotype* interaction terms between environmental conditions are shown. In panels A, a 2-way ANOVA with Tukey’s HSD test was performed. In panels C and D, a Scheirer Ray Hare test and Dunn’s post hoc test with a Benjamini-Hochberg correction was used. Black letters indicate statistically significant means (p< 0.05) within each genotype. Red letters indicate statistically significant means (p< 0.05) across the whole experiment.

This finding prompted us to look at the root elongation phenotypes in our GFP-HY5 over-expressor lines. We found that *GFP-HY5 / hy5-2* had a similar phenotype to the wild type, with no temperature-induced root elongation in the absence of sorbitol, but with temperature-induced root elongation in the presence of sorbitol (Figure 4D). The *snrk2*.*2/snrk2*.*3* mutant behaved as previously, with no temperature-induced root elongation in either the presence or absence of sorbitol. Interestingly though, the overexpression of *GFP-HY5* in this mutant background was able to rescue the phenotype (Figure 4D). This suggests that SnRK2.2 and SnRK2.3 may be a pre-requisite for basal HY5 stability in the root, but they are not *per se* required for changes in HY5 abundance in response to warm temperature and mild osmotic stress.

Together, our findings lead us to propose a model whereby the stability of HY5 in hair cells of the elongation zone is increased by warm temperature and mild osmotic stress. This stabilisation of HY5 correlates with an increase in root elongation in these conditions. In the absence of mild osmotic stress, COP1 promotes the degradation of HY5 specifically in these cells. In the presence of mild osmotic stress, COP1 abundance is reduced, and this allows for the stabilisation of HY5. We propose that SnRK2.2 / SnRK2.3 are required to maintain a background pool of HY5, and as such, warm temperature and osmotic stress no longer have a combined effect on root elongation in the *snrk2*.*2/ snk2*.*3* mutant (Figure S4C).

The evidence presented in this manuscript paints a picture of a complex regulatory network that controls root elongation in the presence of two commonly co-occurring stresses. It is possible that this network operates as a negative feedback loop. HY5 directly promotes the expression of *COP1*^18^, and so our unexpected finding of low *pCOP1:mCherry-COP1* abundance in the *snrk2*.*2 / snrk2*.*3* background (Figure S3D-E) may be due to low HY5 stability in this line. We are not certain about how our data on the *6xABRE_A:erGFP* reporter should be interpreted (Figure 2B). It is possible that this reporter is not reporting on SnRK2 activity *per se*, but rather on HY5 activity in the whole meristem. The *6xABRE_A* reporter contains multiple repeats of an ABA RESPONSIVE ELEMENT from the *ABI1* promoter^9^. HY5 also binds to the promoter of this gene^19^ and so the activation of this reporter may reflect the general increase in HY5 stability seen at warm temperatures (Figure S4A-B).

Our study raises several important questions regarding the molecular basis for mild osmotic stress and warm temperature signalling in the root. Firstly, what is the basis for HY5 stabilisation at warm temperature? We have shown that COP1 and SnRK2.2/ SnRK2.3 gate this response, but we do not know what factor actually promotes HY5 stability in these conditions. Plant shoots contain a multitude of temperature sensors and so it is feasible that multiple points of temperature signal input also exist in roots. Secondly, why do plant roots only elongate at warm temperature in the presence of osmotic stress? We present genetic evidence for the role of *COP1* in controlling this process (Figure 3A), but the reduction in mCherry-COP1 accumulation at 28°C in the absence of sorbitol suggests additional signals are required to activate HY5 in water stress conditions.

This study establishes that in plant roots, warm temperature and water stress signalling are intimately entwined. This finding makes sense on a physiological level, as these environmental factors often occur in combination^20^. Importantly, although we approached this study from the point of view of temperature signalling, our results can also be viewed from the opposite perspective. Warm temperature enhances the response of roots to water stress. It may be that the increased water demands of the shoot at warm temperature^1^ necessitate that warm temperature and water stress signalling are tightly integrated in roots. Our results provide a valuable starting point for further investigations into the combined control of root elongation by warm temperature and mild osmotic stress. We hope that future research leads to further elucidation of the molecular basis and physiological relevance of this response.

## Materials and Methods

### KEY RESOURCES TABLE

See Appendix 1

### CONTACT FOR REAGENT AND RESOURCE SHARING

Further information and requests for resources and reagents should be directed to and will be fulfilled by the Lead Contact, Christa Testerink

### EXPERIMENTAL MODEL AND SUBJECT DETAILS

#### Pre-existing lines

The main experimental organism used in this study was *Arabidopsis thaliana*. Several mutant Arabidopsis lines were described previously: *cop1-4*^13^, *cop1-6*^13^, *snrk2*.*2*/*snrk2*.*3*^8^, *6xABRE-A:erGFP*^9^, *abaQ*^21^, *arebQ*^22^, *cop1-4*/*hy5-2*^23^, *pifq / pif7*^24^, *phyAB*^25^ and *p35s:GFP-HY5 / hy5-2*^17^ are in the Col-0 background. The *abi1-1*^26^ mutant is in the L*er* background. The original sources for these lines can be found in the Key Resources Table.

In addition to Arabidopsis, lettuce (*Lactuca sativa* “Hilde II”), leek (*Allium porrum* “Herfstreuze 2”) and radish (*Raphanus sativus* “Saxa 2”) were used.

#### Lines developed in this study

##### pCOP1:mCherry-COP1

A COP1 promoter fragment was PCR amplified from Col-0 genomic DNA using primers ah930/ah931, digested with SpeI and SbfI, and ligated into *pPPO30A-phyA* ^27^ cut with AvrII/SbfI to replace the *pPHYA:PHYA-YFP-terRbcS* cassette and resulting in plasmid *pPPO-pCOP1 (*#3691). The RbcS terminator was then cut from *pCHF70HA* ^28^ using XbaI/SbfI and ligated into *pPPO-pCOP1 (*#3691) in the XbaI/SbfI sites, resulting in *pPPO-pCOP1-BglII-XbaI-terRbcS* (#3694). Next, *myc-mCherry* CDS was PCR amplified from *pCHF150myc* ^28^ (#2958) using primers ah933/ah934, digested with BamHI/XbaI, and ligated into *pPPO-pCOP1-BglII-XbaI-terRbcS* (#3694) after cutting with BglII/XbaI. This resulted in plasmid *pPPO-pCOP1-myc-mCherry-BglII-XbaI-RbcSter (*#3702). Finally, COP1 CDS was PCR amplified from *pPPO70v1HA-COP1* ^29^ (#3100) using primers ah935/ah227, cut with BamHI/SpeI, and ligated into *pPPO-pCOP1-myc-mCherry-BglII-XbaI-RbcSter* (#3702) after digestion with BglII/XbaI. This resulted in pPPO-pCOP1-myc-mCherry-COP1 (#3749). Plasmid #3749 *pPPO-pCOP1-myc-mCherry-COP1* was transformed into Agrobacteria (*Rhizobium radiobacter*) by electroporation. Agrobacteria (C58) were then used for transformation of Col-0 plants by floral dip^30^. Transgenic lines were selected using 7.6 μl l-1 Inspire (Syngenta Agro AG, Dielsdorf, Switzerland)^27^. Homozygous lines were confirmed by screening seedlings for expression of myc-mCherry-COP1 using epifluorescence microscopy. A map of the final plasmid used for transformation will be made available along with the raw data of this manuscript.

##### pCOP1:mCherry-COP1 / snrk2.2 / snrk2.3

This line was generated through crossing *pCOP1:mCherry-COP1* and *snrk2*.*2* / *snrk2*.*3*. F2 Lines were initially screened through their ability to germinate on 30 nM ABA plates. Homozygous lines were confirmed by PCR.

##### cop1-4/snrk2.2/snrk2.3

This line was generated through crossing *cop1-4* and *snrk2*.*2* / *snrk2*.*3*. F2 Lines were initially screened through their ability to germinate on 30 nM ABA plates, and a *cop1* mutant phenotype in the dark. Homozygous lines were confirmed by PCR.

##### p35s:GFP-HY5 / hy5-2 / snrk2.2 / snrk2.3

This line was generated through crossing *p35s:GFP-HY5 / hy5-2* and *snrk2*.*2* / *snrk2*.*3*. F2 Lines were initially screened through their ability to germinate on 30 nM ABA plates, and the detection of GFP signal in the roots. Homozygous lines were confirmed by PCR.

##### p35s:GFP-HY5 / hy5-2 / cop1-4

This line was generated through crossing *p35s:GFP-HY5 / hy5-2* and *cop1-4*. F2 Lines were initially screened through their *cop1* phenotype in the dark, and the detection of GFP signal in the roots. Homozygous lines were confirmed by PCR.

### PLANT PROPAGATION

To generate the Arabidopsis seed used in this study, plants were grown in rockwool substrate in greenhouses at a 16-hour photoperiod and a temperature of 21°C. Nutrients were supplied throughout the growth cycle through Hyponex solution (pH5.8). Plants were kept well-watered until silique senescence, after which water was increasingly withheld until the plant was fully senesced. Harvested seeds were stored dry at room temperature for at least 2 weeks before use.

### METHOD DETAILS

#### Root and Hypocotyl Length assay

For physiological assays of Arabidopsis plants conducted in soil, seeds were sown directly onto 1kg of soil at either 13%, 15%, 17%, 19%, 21% or 23% water content. Seeds were then germinated for 3 days under a propagator lid (150μE white light, 16h photoperiod, 20°C). No noticeable difference in germination time was observed in these conditions. At day 3, propagator lids were removed, and pots were then placed at either 20°C or 28°C with the same lighting conditions for a further 4 days. Soil water content was re-established daily by dripping water onto the soil surface to a weight of 1kg. On day 7, seedlings were excavated. This was achieved through gently holding them under the cotyledons, whilst water was sprayed to excavate the root. Seedlings were then laid onto agar plates and scanned.

For physiological assay of Arabidopsis plants conducted on agar plates, seeds were surface-sterilised and then sowed on 1/10^th^ Murashige and Skoog medium including vitamins (Duchefa Biochemie), supplemented with 0.5 g/ l MES (Duchefa Biochemie) and 0.8% Diashin Agar (Duchefa Biochemie), pH5.8. After 3-4 days of stratification in dark at 4°C, plates with seed were then placed vertically at 90° in a growth chamber for 3 days in white light (100μE, 16h photoperiod, 20°C) to stimulate germination. At day 3, roots were detached and placed on the same media, supplemented with 5 mM sucrose (Duchefa Biochemie) unless otherwise stated. For osmotic stress and ABA treatments, the medium was additionally supplemented with 25 mM sorbitol (Duchefa Biochemie), 25 mM mannitol (Duchefa Biochemie), 12.5 mM NaCl (Duchefa Biochemie) or 30 nM ABA (Duchefa Biochemie) unless otherwise stated. Root tip position was marked after detachment, and roots were grown in darkness for another 4 days at either 20°C or 28°C before being measured.

For physiological assay of lettuce and radish and leek, plants were grown as stated above for Arabidopsis, but with an altered timetable. Lettuce and radish were germinated for 4 days in white light, and detached roots were grown for another 4 days with or without sorbitol, and at either 20°C or 28°C. Leek seeds were germinated for 7 days at the same conditions, and detached roots were grown for a further 4 days before plates were scanned.

### Abscisic acid level measurements

Wild type Arabidopsis (Col-0) were grown on plates similarly to as above for root lengths assays. One minor modification was that seeds were germinated on plates covered with sterile mesh strips to facilitate the dissection. 60 roots (approximately 5 mg) per sample were harvested in 2ml Eppendorf tubes containing two 1/8” steel ball bearings (Weldtite, 3906141) and flash frozen in liquid nitrogen. Extraction and purification was performed as previously described ^31^, with minor modifications. Samples were ground 2 x 30s at 25 rev/s. Stable isotope-labeled internal standards (100 nM in 10% methanol) were added to ground samples (see Supplementary Table S2). Solvents were removed with a speed vacuum system (thermoSavant) and a StrataX 30mg/3ml spe-column (Phenomenex) was used for purification.

ABA detection and quantification was done using liquid chromatography-tandem mass spectroscopy ^32^. Sample residues were dissolved in 100μL acetonitrile /water (20:80 v/v) and filtered through a 0.2 μm nylon centrifuge spin filter (BGB Analytik). Retention time was assessed using a Waters XevoTQS mass spectrometer equipped with an electrospray ionization source coupled to an Acquity UPLC system (Waters). Acetonitrile/water (+ 0.1 % formic acid) on a Acquity UPLC BEH C18 column (2.1 mm x100 mm, 1.7μm, Waters) at 40 °C with a flowrate of 0.25 mL/min was used to perform chromatographic separations. The column was equilibrated for 30 minutes with the solvent (acetonitrile /water (20:80 v/v) + 0.1% formic acid). 5 μL of sample was injected for analysis, followed by an elution program where the acetonitrile fraction linearly increased from 20% (v/v) to 70% (v/v) in 17 minutes. The acetonitrile fraction was increased between samples to 100% and maintained there for one minute to wash the column. The acetonitrile fraction was set to 20 % before injecting the next sample in one minute and maintained at this concentration for one minute. A capillary voltage of 2.5 kV was combined with a source temperature of 150 °C and desolvation temperature of 500 °C. Quantification was done using multiple reaction monitoring. MRM-setting optimization for the different compounds was done using the IntelliStart MS Console (Supplementary Table S2). Peaks were analyzed using Targetlynx software and samples were normalized for the internal standard recovery (ABA) and expressed relative to the sample fresh weight. Concentration (pmol/mg fresh weight) was determined using a standard curve.

### Confocal imaging

All roots were fixed in 4% paraformaldehyde (Sigma) dissolved in phosphate-buffered saline (PBS, Merck) for 30 minutes under a vacuum, before washing twice in PBS. In some cases, tissues were cleared using ClearSee^33^ (10% w/v xylitol (Sigma), 15% w/v sodium deoxycholate (Sigma), 25% w/v urea (Sigma) for between 1 day and three weeks depending on the experiment. Cell walls were stained with 0.05% Calcofluor white (Megazyme) in ClearSee for 30min and then destained in ClearSee for 30min. Images were collected using a Leica TCS SP8 HyD confocal microscope. For Calcofluor white, an excitation laser of 405 nm and a 425 nm to 475nm band-pass filter was used. For mCherry, an excitation laser of 552 nm and a 590 nm to 630 nm band-pass filter was used. For GFP, an excitation laser of 448 nm and 500 nm to 550 nm band-pass filter was used. Images were acquired using a HC PL APO CS2 63x NA1.40 oil immersion or a 40x water immersion objective fitted with a HyD detector. Within experiments, pinhole, gain, laser power, mode of detection, dynamic range and detector offset were kept constant. Orthogonal sections were re-constructed from z-stack images within the Leica Las-X imaging software.

## QUANTIFICATION AND STATISTICAL ANALYSES

### Image quantification

For root length data, plates were scanned on a flatbed scanner. Scans was quantified in ImageJ, using the freehand line tool.

For fluorescence intensity measurements, confocal images in .lif format were opened in ImageJ. In figure 2, sum stack Z-projections were made, and the region of interest (either the QC or CSC) was highlighted and mean grey value collected. In figure 4, nuclei were first isolated by thresholding before selection and quantification.

### Data presentation

Individual figures were made in R, using the “ggpubr” package. Final figures were collated in Inkscape.

### Statistical analyses

Statistical tests were performed in R. Each figure in the manuscript represents a single experiment. Each experiment was performed at least twice with similar results. For each individual experiment, a Levene test (“car” package) was performed to test for homogeneity of the residuals. For datasets that had equal variance, a 2-way ANOVA (“stats” and “multcompView” packages) was performed. For datasets in which the variance was not homogenous, a Scheirer Ray Hare test (“rcompanion” and “FSA” packages) was performed.

## Supporting information

Key Resources Table

## ACKNOWLEDGEMENTS

Work in Wageningen was supported by grants from European Research Council (ERC) under the European Union’s Horizon 2020 research and innovation programme (grant agreement no. 724321; ERC Consolidator Grant Sense2SurviveSalt-CT), The Dutch Research Council (NWO) (Vici grant; VI.C.192.033-CT) and the Wageningen Graduate School Postdoc Talent Programme (2019-SH).

Work in Freiburg was supported by the German Research Foundation (DFG) under Germany’s Excellence Strategy (EXC-2189 – Project ID 390939984-AH).

We would like to thank Ronald Pierik (Wageningen University) for supplying the *snrk2*.*2/2*.*3* line, Salomé Prat (CRAG, Barcelona) for the *cop1-4, cop1-6, pifqpif7, abaQ, arebQ* and *phyAB* lines, Charlotte Gommers (Wageningen University) for the *cop1-4* / *hy5-215* line, José Dinneny (Standford University) for the *6xABRE-A:erGFP* line and Sang Yeol Lee (Gyeongsang National University) for the *p35S:GFP-HY5 / hy5-2* line. This work would not have been possible without your generosity.

## DATA AVAILABILITY

All raw data and the code used to analyse and present this data will be posted at an online repository upon publication of this manuscript. For access to biological stocks, please get in touch with the Lead Contact, Christa Testerink at.

---

**Figure S1.**
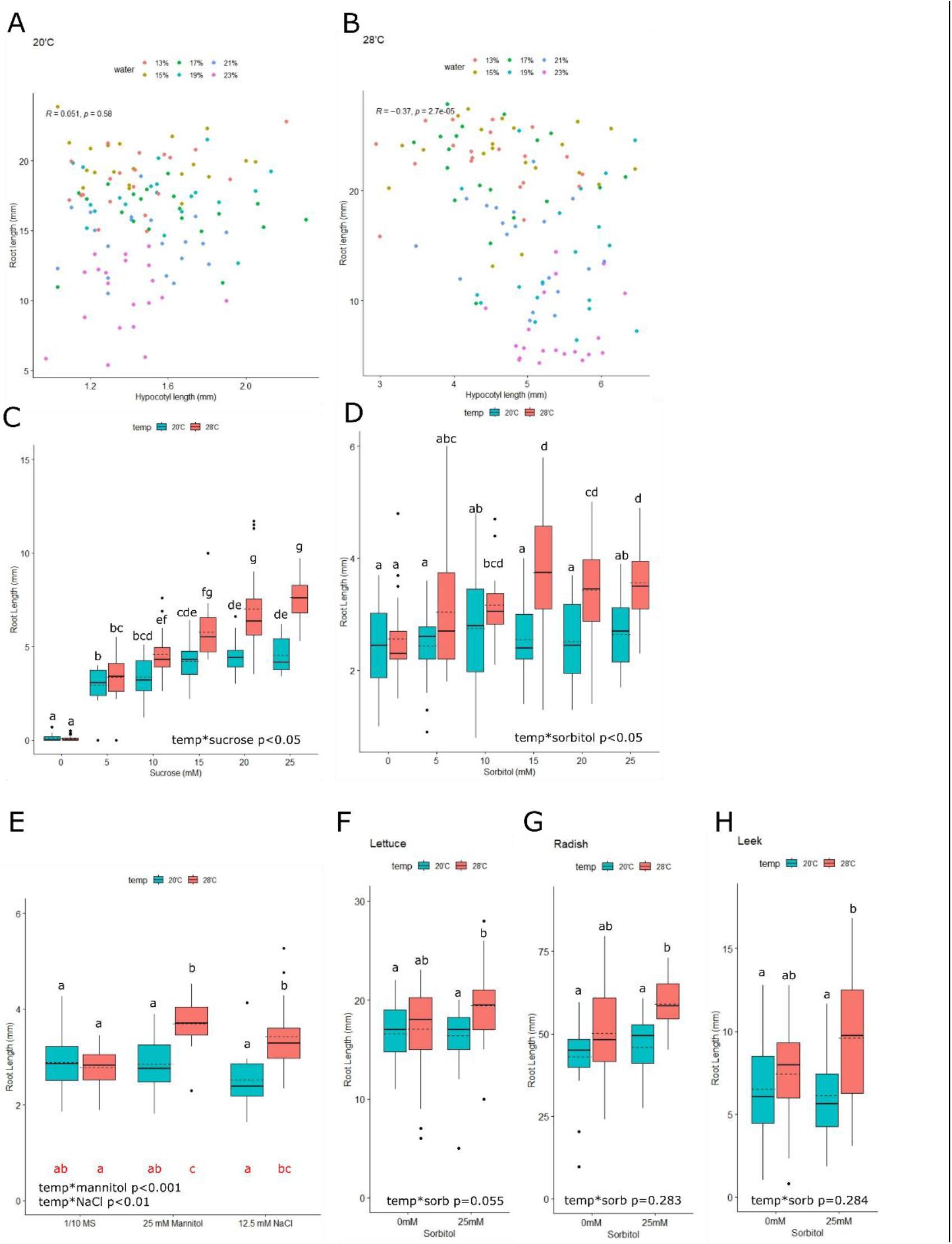
Warm temperature and mild water stress co-operatively promote root elongation. **A)** Correlation between root length and hypocotyl length of wild type (Col) Arabidopsis plants grown on soil at different water availabilities. Plants were sown directly onto soil of the indicated water content and germinated for 3 days under a propagator lid (150μE white light, 16h photoperiod, 20°C). Lids were removed, and pots were treated at 20°C with the same lighting conditions for a further 4 days. Soil water content was re-established daily (n ≥ 19). **B)** Plants grown as in A, but for 4 days at 28°C after removal of the propagator lid. Note that panel S1A and S1B are a reanalysis of the data presented in Figure 1A and Figure 1B (n ≥ 19). **C)** Root length of Col seedlings that were germinated for 3 days in white light (150μE, 16h photoperiod, 20°C) on 1/10^th^ MS media, before roots were detached and placed on 1/10^th^ MS media supplemented with various concentrations of sucrose. Roots were grown in darkness for another 4 days at either 20°C or 28°C before measurement (n=24). **D)** Root lengths of Col seedlings that were germinated for 3 days in white light (150μE, 16h photoperiod, 20°C) on 1/10^th^ MS media, before roots were detached and placed on 1/10^th^ MS media supplemented with 5mM sucrose and various concentrations of sorbitol. Roots were grown in darkness for another 4 days at either 20°C or 28°C before measurement (n=24). **E)** Root lengths of lettuce seedlings that were germinated for 4 days in white light (150μE, 16h photoperiod, 20°C) on 1/10^th^ MS media, before roots were detached and placed on 1/10^th^ MS media supplemented with 5mM sucrose and 0 or 25mM sorbitol. Roots were grown in darkness for another 4 days at either 20°C or 28°C before measurement (n = 28). **F)** Root lengths of radish seedlings that were germinated for 4 days in white light (150μE, 16h photoperiod, 20°C) on 1/10^th^ MS media, before roots were detached and placed on 1/10^th^ MS media supplemented with 5mM sucrose and 0 or 25mM sorbitol. Roots were grown in darkness for another 3 days at either 20°C or 28°C before measurement (n ≥ 15). **G)** Root lengths of leek seedlings that were germinated for 7 days in white light (150μE, 16h photoperiod, 20°C) on 1/10^th^ MS media, before roots were detached and placed on 1/10^th^ MS media supplemented with 5mM sucrose and 0 or 25mM sorbitol. Roots were grown in darkness for another 3 days at either 20°C or 28°C before measurement (n ≥ 24). **H)** Root lengths of Col and *snrk2*.*2* / *snrk2*.*3* seedlings that were germinated for 3 days in white light (150μE, 16h photoperiod, 20°C) on 1/10^th^ MS media, before roots were detached and placed on 1/10^th^ MS media supplemented with 5 mM sucrose and control, mannitol of NaCl (n=16). Roots were grown in darkness for another 4 days at either 20°C or 28°C before measurement. In panels C-G, interaction terms between environmental conditions are shown. In panels A and B, Spearman’s rank correlation coefficient was performed. In panels C and D, a Scheirer Ray Hare test and Dunn’s post hoc test with a Benjamini-Hochberg correction were used. In panels F-G a 2-way ANOVA with Tukey’s HSD test was used. Black letters or asterisks indicate statistically significant means (p< 0.05) within each genotype. Red letters indicate statistically significant means (p< 0.05) across the whole experiment.

**Figure S2.**
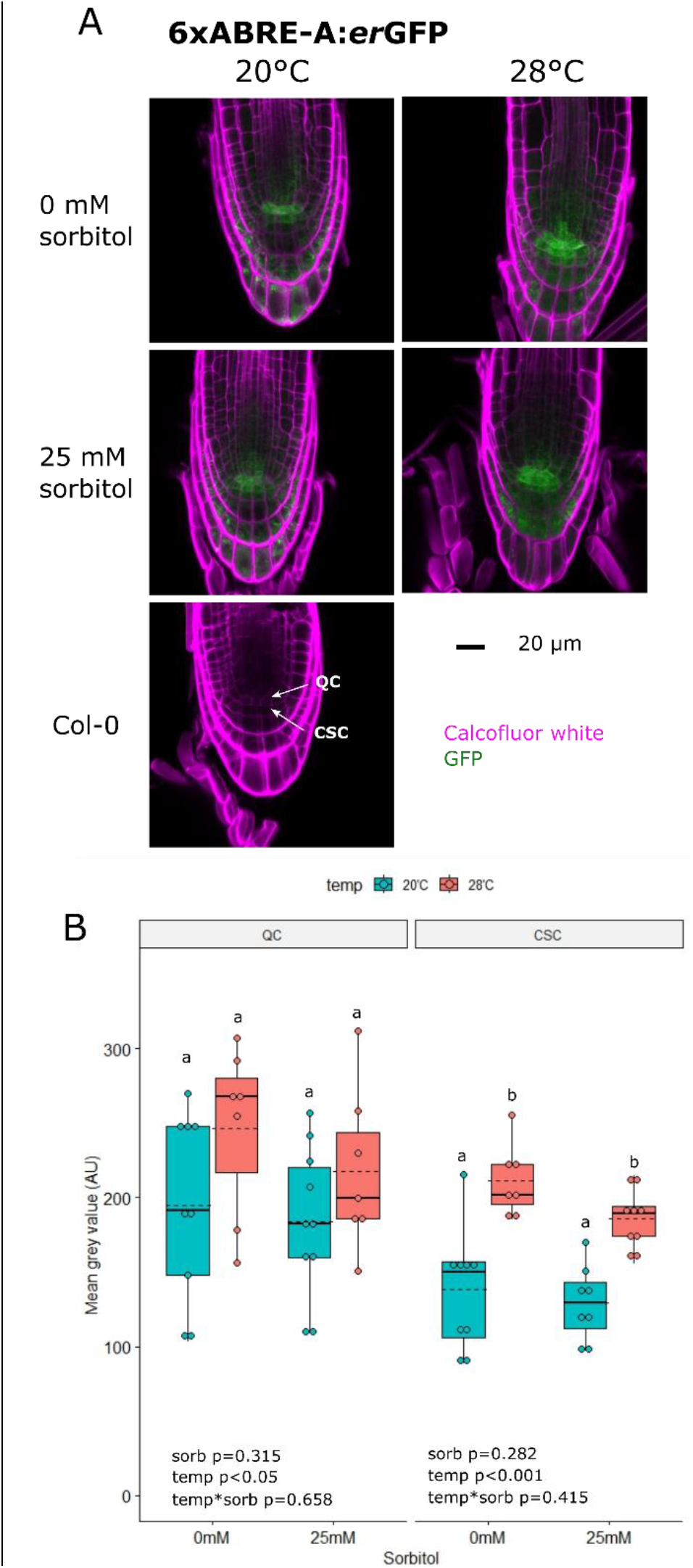
Warm temperature promotes ABA signalling in the root tip. **A)** Uncropped sum-stack images of the root tips shown in Figure 2B. Quiescent centre (QC) and Columella stem cells (CSC) are indicated. **B)** Quantification of GFP signal intensity from the QC and CSC of at least seven plants grown as in Figure 2B (n ≥ 7). For each tissue type, a 2-way ANOVA with Tukey’s HSD test was performed. Interaction terms between environmental conditions are shown. Different letters or asterisks indicate statistically significant means within each tissue.

**Figure S3.**
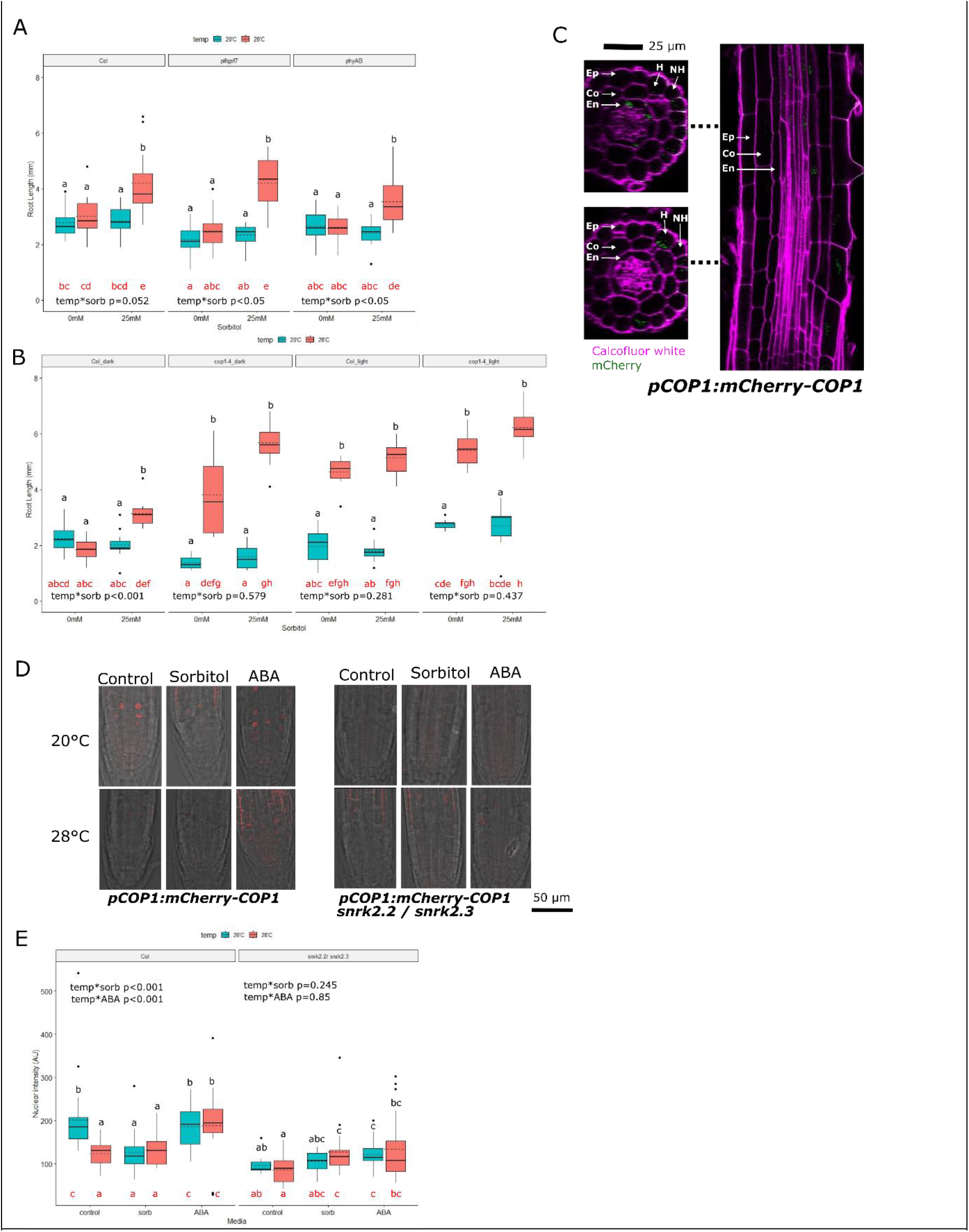
COP1 supresses warm temperature signalling in the absence of osmotic stress. **A)** Wild type Col, *pif1-1 / pif3-3 / pif4-2 / pif5 / pif7-1* (*pifq / pif7*) and *phyA / phyB-9* (*phyAB*) mutant plants were germinated for 3 days in white light (150μE, 16h photoperiod, 20°C) on 1/10th MS media, before roots were detached and placed on 1/10th MS media supplemented with 5 mM sucrose and 0 mM or 25 mM sorbitol. Roots were grown in darkness for another 4 days at either 20°C or 28°C before being measured (n ≥ 14). **B)** Wild type Col or cop1-4 mutant plants were grown as in A, but after detachment, roots were grown for 4 days in either darkness or 150μE white light with a 16h photoperiod (n ≥ 8). **C)** *pCOP1:mCherry-COP1* were grown for three days in white light (150μE, 16h photoperiod, 20°C) on 1/10th MS media, before roots were detached and placed on 1/10th MS media supplemented with 5 mM sucrose and 25 mM sorbitol. Roots were grown in darkness for another 4 days at 20°C before fixation. Virtual cross sections were made through the differentiation zone (upper) and the end of the elongation zone (lower). Epidermis (Ep), cortex (CO), and endodermis (En) are indicated. Within the epidermis, cells are also characterised as either hair (H) or non-hair (NH) cells depending on their number of contacts with cortex cells. **D)** Representative confocal microscopy images of *pCOP1:mCherry-COP1* roots in both the wild type and *snrk2*.*2 / snrk2*.*3* background. for three days in white light (150μE, 16h photoperiod, 20°C) on 1/10th MS media, before roots were detached and placed on 1/10th MS media supplemented with 5 mM sucrose and 25 mM sorbitol, 30 nM ABA or a negative control. Roots were grown in darkness for another 4 days at 20°C before fixation. Images show a 10μm sum stack projection, with red mCherry signal overlayed on a brightfield image in grey. **E)** Quantification of nuclear mCherry signal intensity of the three brightest nuclei from images of roots grown as in D (n ≥ 6 roots). In each quantitative panel, *within genotype* interaction terms between environmental conditions are shown. In all panels, Scheirer Ray Hare test and Dunn’s post hoc test with a Benjamini-Hochberg correction was used. Black letters indicate statistically significant means (p< 0.05) within each genotype / light condition. Red letters indicate statistically significant means (p< 0.05) across the whole experiment.

**Figure S4:**
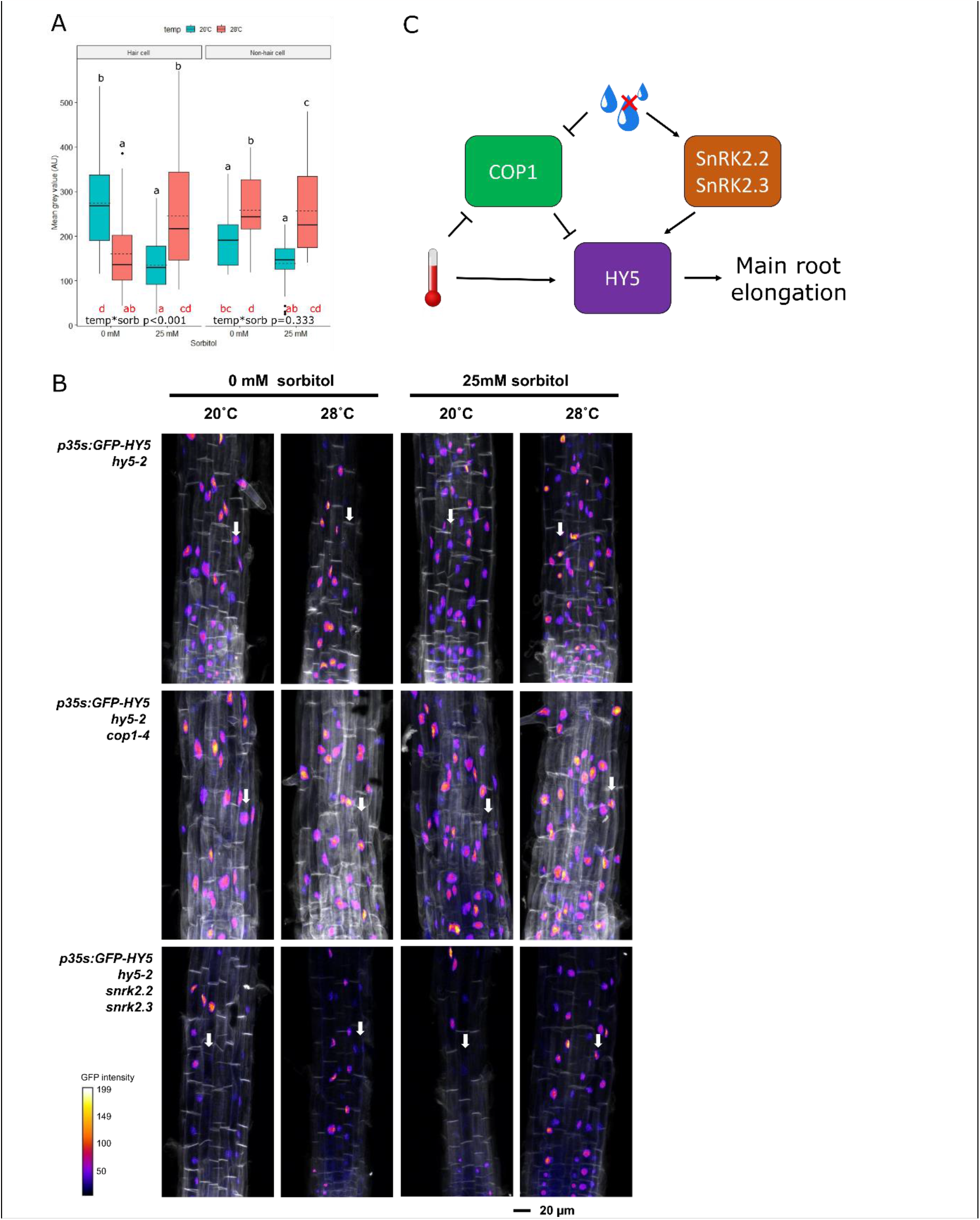
Warm temperature and mild water stress promote the accumulation of HY5. **A)** Mean grey value of hair and non-hair cell nuclei from sum stack images of *p35S:GFP-HY5 / hy5-2* roots. Plants were grown for 3 days in white light (150μE, 16h photoperiod, 20°C) on 1/10^th^ MS media, before roots were detached and placed on 1/10^th^ MS media supplemented with 5 mM sucrose and 0 mM or 25 mM sorbitol. Roots were grown in darkness for another 4 days at either 20°C or 28°C before fixation. A Scheirer Ray Hare test and Dunn’s post hoc test with a Benjamini-Hochberg correction was used to determine interactions between environmental conditions. Black letters indicate statistically significant means (p< 0.05) within each cell type. Red letters indicate statistically significant means (p< 0.05) across the whole experiment. At least three nuclei were quantified for each root (n=6 roots). **B)** Uncropped sum-stack images of the root tips shown in Figure 4B. Hair cell files are indicated with white arrows. C) A proposed working model for co-operative regulation of root elongation by water stress and warm temperature. In the absence of water stress and at 20°C, COP1 supresses HY5 in the elongation zone root hair cells. Under mild water stress at 20°C, COP1 abundance is reduced, but in the absence of warm temperature, this is not enough to promote HY5 stability. At 28°C, COP1 abundance is reduced, but this does not result in HY5 stabilisation if no water stress is present. In the presence of both warm temperatures and mild water stress, COP1 abundance is reduced, and HY5 stability is enhanced (possibly through the action of SnRKs).

